# Measuring American black bear genetic load in fixed structural variants and runs of homozygosity

**DOI:** 10.64898/2026.11.05.764578

**Authors:** Faezeh Azimi Chetabi, Emily E. Puckett, Joseph M. Northrup, Aaron B. A. Shafer

## Abstract

Populations living in geographically isolated regions are expected to have reduced gene flow, increased genetic drift, and elevated levels of inbreeding, influencing both genetic load and possibly the distribution of structural variants. The American black bear (*Ursus americanus*) is widely found across Ontario, Canada, where populations inhabit diverse landscapes ranging from relatively intact contiguous forested landscapes to isolated regions such as the Saugeen Peninsula. Here, we investigated the population genomic history across five Ontario black bear populations using a combination of short-read (SR) and long-read (LR) sequencing data. We compared population genetic metrics including nucleotide diversity (π), Tajima’s D, runs of homozygosity (ROH), and genome-wide structural variants (SVs) to assess population health and identify signatures of isolation. Our results showed that the isolated Saugeen Peninsula population is genetically distinct and exhibited reduced nucleotide diversity, greater ROH and SV abundance compared with mainland populations. The population also showed elevated genetic load in all genomic features, suggesting increased exposure of deleterious variation that are expressed in ROH and population specific SVs. We further discuss the integration of ROH, SVs and genetic load in conservation assessments.

## 1. Introduction

Population isolation represents a significant conservation challenge because reduced gene flow into small and fragmented populations promotes genetic drift and inbreeding (Schlaepfer et al., 2018). These processes diminish genetic diversity and increase the likelihood of long-term population decline (Draper et al., 2017; Clendenin et al., 2025). Habitat fragmentation, geographic barriers, and human infrastructure can restrict dispersal among populations, thereby reducing genetic connectivity even before pronounced genetic differentiation becomes evident (Proctor et al., 2020; Wilton et al., 2015). For example, isolated Arctic fox (*Vulpes lagopus*) populations tend to exhibit increased runs of homozygosity (ROH), elevated genetic load, and reduced overall genetic diversity which may lead to expression of harmful and negative effect on individual fitness and survival (Hasselgren et al., 2024). While purging appears relatively common (Wootton et al, 2023), long-term isolation does not necessarily result in the purging of deleterious genetic variation. In beluga whale populations (*Delphinapterus leucas*), sustained isolation over hundreds of generations was associated with the accumulation of deleterious mutations that has led to a high mutation load without evidence of effective genetic purging, suggesting that specific demographic histories can strongly influence the genomic consequences of isolation (Orton et al., 2026).

Single nucleotide polymorphisms (SNPs) are a staple of conservation genomics due to their abundance throughout the genome and their ability to provide high-resolution insights into population genetic processes. Compared with smaller marker panels, large SNP datasets can improve the resolution of population structure, individual assignment, dispersal inference, admixture, and phylogeographic history (Miller et al., 2012; Puckett et al., 2015; Puckett and Eggert, 2015; Petrou et al., 2025). However, SNPs represent only one class of genomic variation and may not fully capture larger-scale genomic changes that influence phenotypes, adaptation, and evolutionary potential. Structural variants (SVs), including insertions, deletions, duplications, inversions, and copy-number changes, encompass substantially larger genomic regions and may alter gene content and order, gene regulation, and trait-associated regions (Wold et al, 2021). Comparative genomic studies have shown that SVs, often combined with SNPs and gene family analysis, can identify candidate loci associated with local adaptation and other biologically important traits (Derežanin et al., 2022; Palma-Vera et al., 2022, Azimi Chetabi & Shafer, 2026).

Structural variants can capture functional genomic variation that is undetected by SNPs alone and therefore improve the assessment of genetic diversity and population health (Wold et al., 2021; Stuart et al., 2025). For example, in the endangered and inbred Hihi (*Notiomystis cincta*; New Zealand stitch bird), SVs with potentially deleterious effects were associated with lifetime fitness and provided information complementary to conventional SNP-based analyses, demonstrating that reliance on SNP data alone might miss important components of genomic variation relevant to conservation (Stuart et al., 2025). Similarly, work on Scandinavian wolves (*Canis lupus*) found that the realized genetic load of SVs increased with inbreeding, whereas immigration reduced this load by restoring genetic diversity and reintroducing ancestral alleles, highlighting the importance of gene flow for mitigating negative genetic consequences (Smeds et al., 2024). Structural variants have also been implicated in inbreeding depression through the accumulation of deleterious mutations within chromosomal inversions, emphasizing the need to consider these variants when developing conservation and genetic management strategies (Schneller et al., 2025). More broadly, the integration of SV datasets has been shown to improve the delineation of conservation units and population structure in endangered species, providing additional information for conservation planning and management (Liu et al., 2026; Wold et al., 2021).

American black bears (*Ursus americanus*) once ranged over much of North America, but habitat destruction and fragmentation associated with European settlement and expansion resulted in drastic range reduction and, in some cases, isolation of populations (Scheik and McCown 2014). Isolated populations of black bears consistently exhibit lower genetic diversity, greater genetic differentiation, and stronger signatures of inbreeding than populations connected through immigration (Belant et al., 2005; Wilton et al., 2015; Draper et al., 2017). Both demographic bottlenecks and founder events have reduced genetic diversity and accelerated genetic drift, while admixture and translocations partially restored genetic variation in some black bear populations (Puckett et al., 2014). These patterns are also evident in black bears found in Ontario, Canada, where most populations remain genetically connected through gene flow across the province, though fine-scale patterns of gene flow are influenced by population density and broad gradients in food productivity (Kopsala et al., 2023). The isolated Saugeen Peninsula (formerly Bruce) population in Ontario forms a genetically distinct cluster maintained by restricted gene flow with mainland populations (Pelletier et al., 2012). Pelletier et al. (2012) showed reduced genetic diversity and limited immigration from surrounding mainland populations, likely reflecting the combined effects of historical isolation, habitat fragmentation, and increasing human development along the southern portion of the peninsula. Although Ontario black bear populations generally exhibit high genetic diversity and connectivity, geographically isolated populations may experience different demographic and genetic processes. In this study, we investigated the genomic consequences of population isolation in Ontario black bears, with a focus on the geographically distinct Saugeen Peninsula (SP) population (Figure 1). We combined analyses of ROHs, SVs, and population genomic metrics to characterize patterns of genetic diversity, inbreeding, and genetic load across Ontario black bear populations.

**Figure 1.**
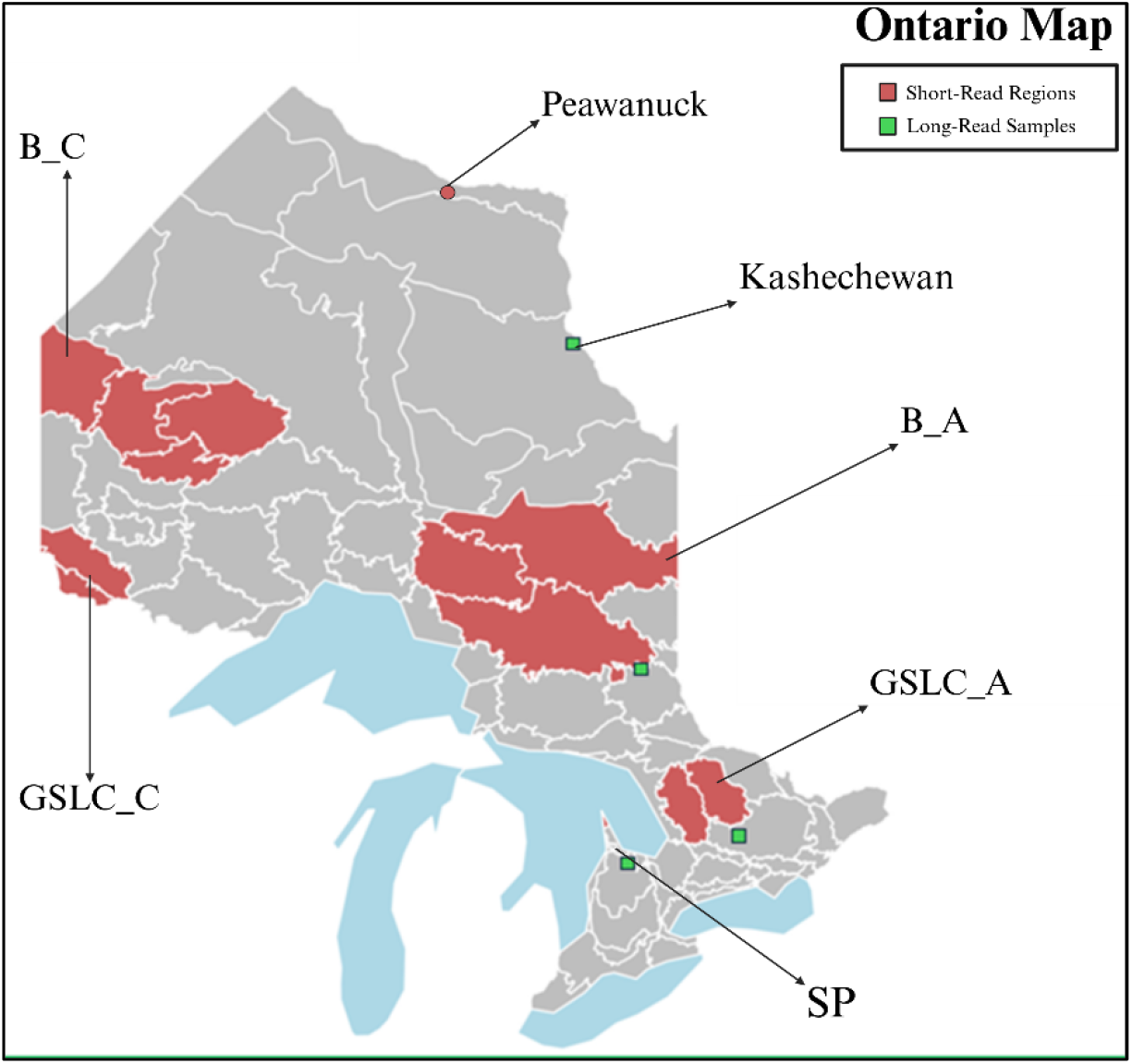
Geographic distribution of sampled Ontario black bear populations. Red polygons highlight the five focal geographic regions where short read WGS was produced from; west to east these include boreal continental (B_C), Great Lakes Saint Lawrence continental (GLSL_C), boreal Atlantic (B_A), Great Lakes Saint Lawrence Atlantic (GLSL_A) and SP (Saugeen Peninsula). We also highlight one individual with a short-read genome from Peawanuck in the far north of the province. Green squares indicate the locations for long read sample from Kashechewan First Nation.

## 2. Materials and Methods

### 2.1. Data gathering and preparation

Bear tooth samples were provided to the Ontario Ministry of Natural Resources through the tooth submission program through which hunters are asked to voluntarily submit teeth from harvested animals. Typically, there is blood or tissue still attached to teeth from which DNA can be extracted. A subset of these were selected for short-read sequencing. We also collected tissue from freshly harvested animals and animals handled as part of ongoing research and management activities in the province. All samples were frozen upon arrival at the laboratory at −20°C until processing. For Illumina short-read (SR) sequencing DNA was extracted from samples using the QIAGEN DNeasy Blood & Tissue Kit, where PacBio long-read (LR) samples were extracted using the Qiagen HMW MagAttract kit. We used the black bear reference genome assembly (JANIGQ000000000) from the National Center for Biotechnology Information (NCBI) database (Supple et al, 2024). Assembly completeness was evaluated using BUSCO (Simão et al., 2015), yielding a score of 99.8%.

We generated 28 Illumina whole-genome SR Illumina datasets and 4 PacBio LR datasets from American black bears across Ontario (Figure 1). We assigned samples to 5 habitat-types according to major forest types (boreal versus mixed deciduous, also known as the great-lakes Saint Lawrence forest; Rowe et al. 1972) and climate zones (continental versus Atlantic; see Kopsala 2022) (Figure 1): 1) boreal continental (B_C; 4 samples), 2) Great Lakes Saint Lawrence continental (GLSL_C; 4 samples), 3) boreal Atlantic (B_A; 5 samples), 4) Great Lakes Saint Lawrence Atlantic (GLSL_A; 7 samples) and 5) SP (Saugeen Peninsula; 10 samples). These samples had locations accurate to the management unit only. Samples from Kashechewan First Nation and Peawanuck (both far north Ontario) were considered part of B_A population and had precise location coordinates (Figure 1). More information can be found in Table S1.

### 2.2. Calculating population diversity and admixture

Raw FASTQ files were trimmed using Trimmomatic (Bolger et al., 2014) with a MINLEN threshold of 36 and aligned to the American black bear reference genome using Bowtie2 (Langmead et al., 2012). SAMtools (Danecek et al., 2021) was used to convert, sort, and index the alignment files, and duplicate reads were removed using Picard (Broad Institute, 2018). Single nucleotide polymorphisms were identified using ANGSD (Korneliussen et al, 2014). Genotype likelihoods were estimated under the SAMtools model (-gl 2) and exported in Beagle format (-doGlf 2). SNPs were retained only if they met a significance threshold of 1 × 10 (-SNP_pval 1e-6), with a minimum mapping quality of 20 (-minMapQ 20) and a minimum base quality score of 20 (-minQ 20). Major and minor alleles were inferred directly from the genotype likelihoods (-doMajorMinor 1; -doMaf; -minMaf 0.1). The resulting BCF file was converted to VCF format using BCFtools (Danecek et al., 2021). Major and minor alleles were inferred directly from the sequencing data, and genotype likelihoods were exported in Beagle format.

For the principal component analysis (PCA), we used the SNP panel from the SR data to minimize potential bias arising from differences in sequencing technologies between SR and LR. ANGSD was first run using only the SR BAM files, and the coordinates of the identified SNPs were extracted. These coordinates were then supplied to a second ANGSD run as a predefined sites file (-sites), which included both SR and LR samples, restricting genotype likelihood estimation to the SNP positions identified from the SR data. The resulting genotype likelihoods were exported in Beagle format and analyzed using PCAngsd (Meisner et al., 2018), which estimates individual covariance matrices directly from genotype likelihoods without requiring explicit genotype calling. We ran PCA with and without the SP samples.

We also used NGSadmix (Skotte et al., 2013), which estimates individual ancestry proportions directly from genotype likelihoods without requiring genotype calling. Genotype likelihoods generated in ANGSD were provided in Beagle format as input for NGSadmix. To explore patterns of genetic clustering among individuals, analyses were performed assuming two, three, and four ancestral populations (K = 2, 3, and 4). For each value of K, ancestry proportions were estimated using a maximum-likelihood framework based on genotype likelihoods. The resulting admixture coefficients were used to visualize population structure and assess the degree of genetic differentiation and admixture among Ontario black bear populations.

Population genetic diversity and differentiation were estimated using ANGSD (Korneliussen et al, 2014) and the realSFS module. For each a-priori population, corresponding to the predefined management units (MUs), folded site allele frequency likelihoods were calculated from genotype likelihoods, and the site frequency spectrum (SFS) was estimated using realSFS. The resulting SFS was used as a prior to estimate neutrality statistics and nucleotide diversity. Tajima’s D and nucleotide diversity (π) were calculated both genome-wide and in sliding windows of 50 kb with a step size of 10 kb using thetaStat. Genetic differentiation among populations was assessed using the fixation index (F_St_) implemented in realSFS. Pairwise joint site frequency spectra were first estimated for all population comparisons and subsequently used to calculate weighted genome-wide F_St_ values. To investigate variation in population differentiation across the genome, windowed F_St_ estimates were calculated using 50 kb windows with a 10 kb step size. To quantify isolation by distance (IBD), geographic distances between populations were calculated using their geographic coordinates to create a pairwise geographic distance matrix. A corresponding genetic distance matrix was created using pairwise F_St_ values. The relationship between genetic and geographic distances was quantified using a Mantel test.

### 2.3. Runs of Homozygosity and Structural Variant Detection

Runs of homozygosity were identified using BCFtools (Danecek et al., 2021), using the Hidden Markov Model (HMM) and genotype likelihood information to infer extended homozygous regions across the genome. Individual ROH results were extracted for each sample and filtered to retain only high-confidence ROHs with a quality score ≥ 50 and with an average ROH quality score ≥ 25. The genomic coordinates of filtered ROHs were converted to BED format and intersected with the American black bear gene annotation using BEDTools (Quinlan et al, 2010) to identify genes located within ROH regions.

For the SV detection we followed the approach of Azimi Chetabi & Shafer (2026); in brief, we removed adaptors from the raw fastq files using Trimmomatic (Bolger et al., 2014) with a MINLEN threshold of 36. We employed Bowtie2 (Langmead et al., 2012) to align all samples to the American black bear genome. We used SAMtools (Danecek et al., 2021) to convert, sort, and index the SAM files. To ensure data integrity, duplicate reads were eliminated using Picard (Broad Institute, 2018) with REMOVE_SEQUENCING_DUPLICATES set to TRUE. We then employed three different SV calling tools for SR data: Delly (Rausch et al., 2012), Manta (Chen et al., 2016), and Lumpy (Layer, 2014), all with default settings. Further refinement and comparative analysis of VCFs were performed using BCFtools (Danecek et al., 2021), retaining only SVs with SUPP > 5, representing the number of sequencing reads supporting a given SV. We merged the SV call sets from Delly, Manta, and Lumpy using Jasmine (Kirsche et al, 2023), configured to confirm SVs found in at least two tools.

For the LR samples, we mapped reads to American black bear reference genome using Winnowmap (Jain et al, 2020). The aligned SAM files were then converted to BAM format, sorted, and indexed with SAMtools (Danecek et al., 2021). Structural variants were identified using a combination of Sniffles (Sedlazeck et al., 2018), SVIM (Heller et al, 2019), and NanoVar (Tham et al, 2020). We employed Jasmine (Kirsche et al, 2023) to merge the SV calls into a single VCF file if SVs are found in at least 2 tools. We filtered the merged VCF files based on read support greater than 5. The resulting impact of SVs from both LR and SR analysis was annotated using SnpEff (Cingolani et al., 2012).

### 2.6. Genetic Load, Gene Impact and Clustering

To investigate the relationship between SVs and inbreeding, SV and ROH coordinates were intersected using BEDTools and overlapping regions between SVs and ROHs were identified separately for each individual. Population-level SVs were identified by comparing the SV call sets of all individuals within a population using Jasmine (Kirsche et al, 2023) and only SVs present in every individual (i.e. fixed) of a given population were kept. Similarly, fixed ROHs were identified by converting filtered ROH coordinates to BED format, sorting and merging overlapping intervals within each individual using BEDTools (Quinlan et al, 2010) and performing multi-sample intersection analyses using the BEDTools multiinter function. ROH regions were considered population-level when they were present in all individuals within a population. To focus on biologically meaningful regions, only fixed ROHs exceeding 100 kb in length were retained for downstream analyses.

Genetic load was calculated using functional annotations of SNPs identified from the final ANGSD dataset. The filtered SNP dataset was annotated using SnpEff (Cingolani et al., 2012). Variants carried by each individual were identified based on genotype information and classified according to their predicted SnpEff impact as HIGH. Missense variants were identified based on the missense_variant annotation, while loss-of-function (LOF) variants were identified using the SnpEff LOF annotation. For each individual, the number of variants in each functional category was calculated by retaining sites at which the individual carried at least one non-reference allele. We separated the distribution of genetic load by SNPs found in ROH, SVs, and the rest of the genome. Differences among populations were first compared using an ANOVA; because of variation in feature density, we resampled with replacement the VCF file used by SnpEff into 100 files of 1,000,000 sites and compared differences as above.

## 3. Results

### 3.1. Population metrics

A total of 8,000,244 SNPs were identified using ANGSD after filtering for site depth (100 ≤ DP ≤ 600). Principal component analysis showed clear genetic differentiation between SP and the mainland regions (Figure 2a); when SP individuals were excluded, further differentiation among the populations was observed (Figure 2b). The SP population had the highest Tajima’s D value (0.53) and the lowest nucleotide diversity (π = 0.0011), while the remaining populations showed negative Tajima’s D values, ranging from −0.20 in B_A to −0.008 in B_C. Fixation index (F_St_) values among Ontario black bear populations ranged from 0.050 to 0.190 (Figure S1a). Among the mainland populations, the lowest level of genetic differentiation was observed between B_A and GLSL_C, though all values excluding comparisons with SP were small; in contrast, SP exhibited substantially greater differentiation from all mainland populations, with FST values of >0.15 (Figure S1a). The relationship between genetic and geographic distances was not statistically significant (R² = 8.2 × 10, p = 0.542), showing no evidence of isolation by distance (Figure S1b).

**Figure 2.**
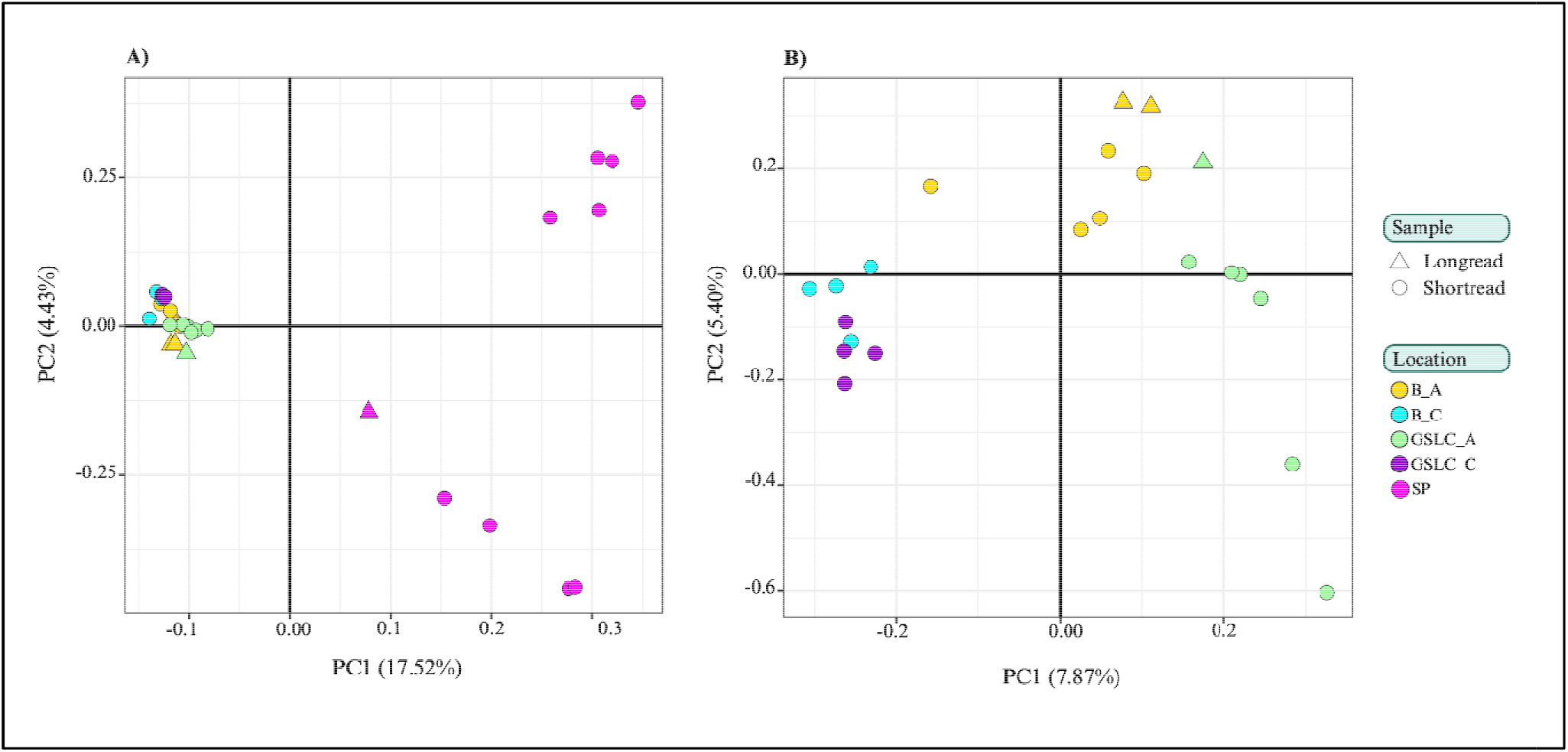
Principal component analysis (PCA) of American black bears across Ontario based on genotype likelihoods using PCAngsd. **a)** PCA including all samples. **b)** PCA excluding samples from Saugeen Peninsula (SP). Individuals are colored according to sampling location, while point shapes indicate sequencing type (long-read or short-read).

NGSadmix analysis with K = 3 identified three major ancestry components among the Ontario black bear populations (Figure S2a). SP was predominantly characterized by a distinct ancestry component (Cluster 3), distinguishing it from the mainland populations, although several SP individuals showed varying contributions from the other ancestry components. Among the mainland populations, B_C and GLSL_C were exclusively assigned to Cluster 2, with both showing little evidence of admixture. In contrast, B_A and GLSL_A exhibited greater admixture between Clusters 1 and 2, with some individuals also showing small contributions from Cluster 3 (Figure S2a, see Figure S2b for K=2).

### 3.2. Inbreeding and Structural Variants

The SP population exhibited the highest F_ROH_ (∼0.25; Table 1) and number of ROH (38,360), though average ROH lengths were similar among populations (Table 2; Figure 3, Figure S4). The SP population had the highest number of fixed ROH shared by all individuals (17), with an average size of 59 kb and a median size of 23 kb (Table 5). We identified a total of 257,699 SVs (Table 2) where deletions represented the majority of variants (60.17%), followed by insertions (39.48%), whereas duplications (0.19%) and inversions (0.16%) were rare. Median variant lengths were 60 bp for deletions, 118 bp for insertions, 1,513 bp for duplications, and 595 bp for inversions. No translocations were identified. The highest number of SVs was detected in the B_A population (106,010), followed by SP (70,911; Table 2). Inversions were detected only in SP (103), B_A (208), and GLSL_A individuals (98; Table 2).

**Figure 3.**
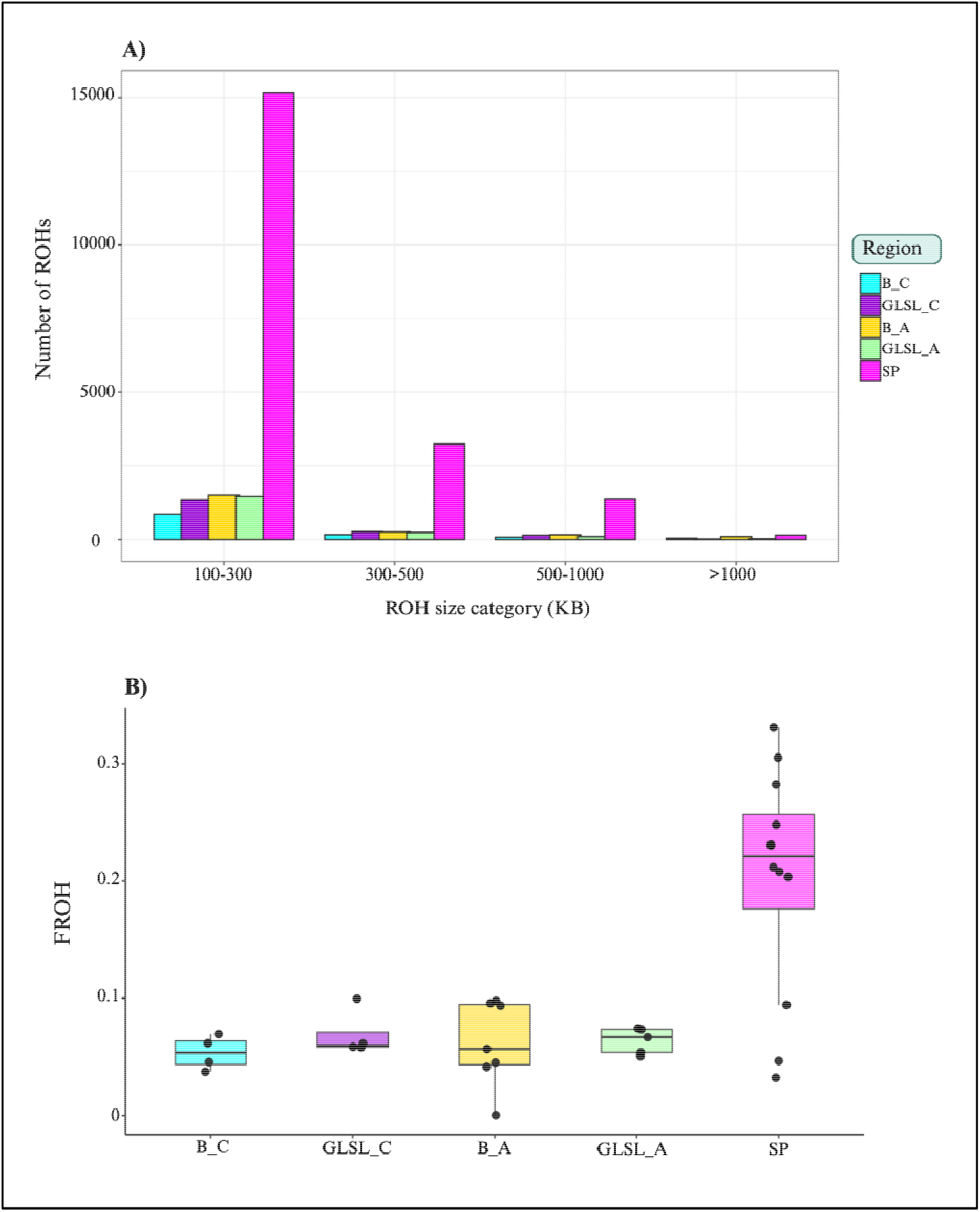
a) Distribution of runs of homozygosity (ROH) across size classes in Ontario American black bear populations. Bars represent the number of ROHs identified in each population across four ROH size categories (100–300, 300–500, 500–1000, and >1000 kb). **b)** Distribution of genomic inbreeding coefficients among Ontario American black bear populations. Boxplots show the distribution of individual FROH values within each population, with black points representing individual bears.

**Table 1.** Summary of genetic diversity and inbreeding metrics among Ontario American black bear populations. Tajima’s D, nucleotide diversity (π), mean genomic inbreeding coefficient (F_ROH_), and the range of average ROH lengths are reported for each population. Sample sizes were SP (n = 10), B_A (n = 5), B_C (n = 4), GLSL_A (n = 7), and GLSL_C (n = 4).

| **Population** | **Tajima’s *D*** | **Nucleotide diversity (π)** | **Mean F<sub>ROH</sub>** | **Range of average ROH length (kb)** |
|---|---|---|---|---|
| **SP** | 0.53 | 0.0011 | 0.25 | 69.96-185.63 |
| **B_A** | -0.2 | 0.0017 | 0.071 | 37.38-111.08 |
| **B_C** | -0.008 | 0.0013 | 0.068 | 61.41-549.11 |
| **GLSL_A** | -0.08 | 0.0015 | 0.075 | 60.25-84.12 |
| **GLSL_C** | -0.03 | 0.0014 | 0.077 | 69.76-105.16 |

**Table 2.**
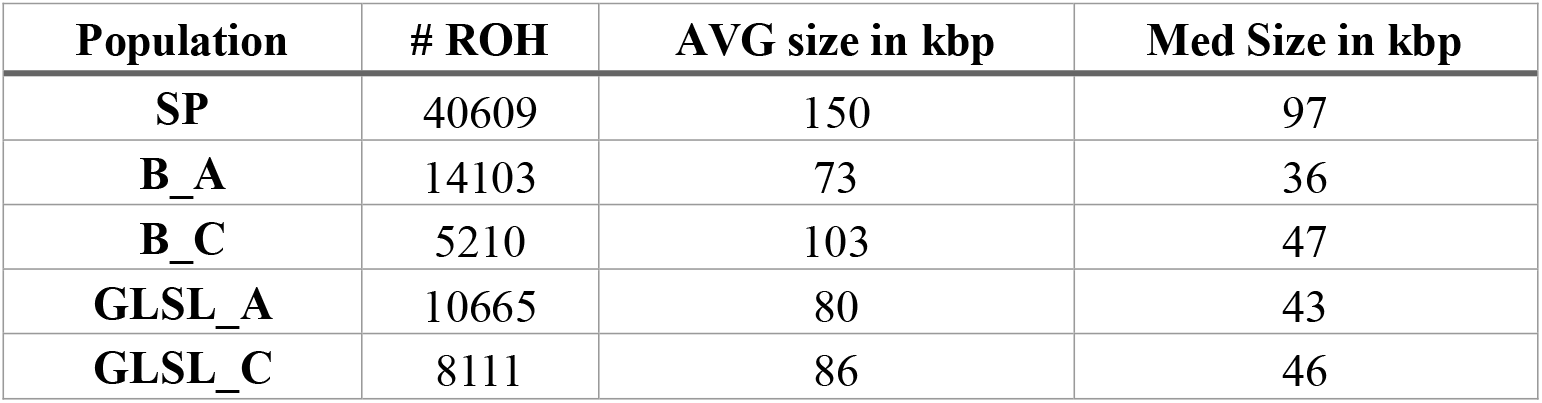
Summary of runs of homozygosity identified among Ontario American black bear populations. The total number of ROHs, average ROH length, and median ROH length are reported for each population.

| **Population** | **# ROH** | **AVG size in kbp** | **Med Size in kbp** |
|---|---|---|---|
| **SP** | 40609 | 150 | 97 |
| **B_A** | 14103 | 73 | 36 |
| **B_C** | 5210 | 103 | 47 |
| **GLSL_A** | 10665 | 80 | 43 |
| **GLSL_C** | 8111 | 86 | 46 |

**Table 3.** Summary of structural variants identified among Ontario American black bear populations. The total number of SVs and the number classified as deletions, duplications, insertions, and inversions are reported for each population.

| **Population** | **# SVs** | **Deletion** | **Duplication** | **Insertion** | **Inversion** |
|---|---|---|---|---|---|
| **SP** | 73873 | 46016 | 192 | 27562 | 103 |
| **B_A** | 106010 | 58549 | 100 | 47153 | 208 |
| **B_C** | 9312 | 7596 | 44 | 1672 | 0 |
| **GLSL_A** | 55621 | 32325 | 78 | 23120 | 98 |
| **GLSL_C** | 12883 | 10568 | 84 | 2231 | 0 |

The number of fixed SVs shared by all individuals within each population ranged from 6 to 28 (Table 4). The SP population contained 11 shared deletions with a median size of 3,160 bp. The B_A population contained 13 shared deletions with a median size of 2,603 bp. Among the remaining populations, GLSL_A contained the highest number of fixed SVs, with 26 deletions and a median size of 664 bp, followed by GLSL_C with 23 deletions and a median size of 486 bp (Table 4). Overall, deletions represented all fixed SVs. The number of SV–ROH overlaps varied among populations, with average per individual slightly elevated in SP but not the highest (Table 6). There were no overlapping SVs and ROHs found in all individuals of a population.

**Table 4.** Summary of fixed structural variants among Ontario American black bear populations. The number of fixed SVs, median SV size, and SV types are reported for each population, including the SP population analyzed with mainland individuals.

| Population | # SVs | Median SV Size | SV type |
|---|---|---|---|
| **SP** | 11 | 3160 | DEL: 11 |
| **B_A** | 13 | 2603 | DEL: 13 |
| **B_C** | 6 | 28 | DEL:4 INS:2 |
| **GLSL_A** | 26 | 664 | DEL: 28 |
| **GLSL_C** | 23 | 486 | DEL: 23 |

### 3.3. Feature Overlap and Genetic Load

The distribution of predicted functional effects of SNPs varied among populations (Figure 4, Figure S3). For genome-wide SNPs after masking ROH and SVs, the SP population exhibited higher overall counts of gene impact variants, particularly MISSENSE and LOF variants, with ANOVA indicating significant differences among populations for HIGH, MISSENSE, and LOF categories (Figure 4a). Among SNPs overlapping ROHs, the SP population again showed the highest counts across all three categories, with differences among populations supported by ANOVA for HIGH, MISSENSE, and LOF variants (Figure 4b). A similar pattern was observed for SNPs overlapping SVs, where SP showed higher counts, particularly for MISSENSE and LOF variants, and ANOVA indicated differences among populations across all three categories (Figure 4c). This same pattern was observed in the bootstrapped data (Figure S3). Among the mainland populations, functional-effect distributions were generally more similar, although variation among populations and individuals was observed (Figure 4).

**Figure 4.**
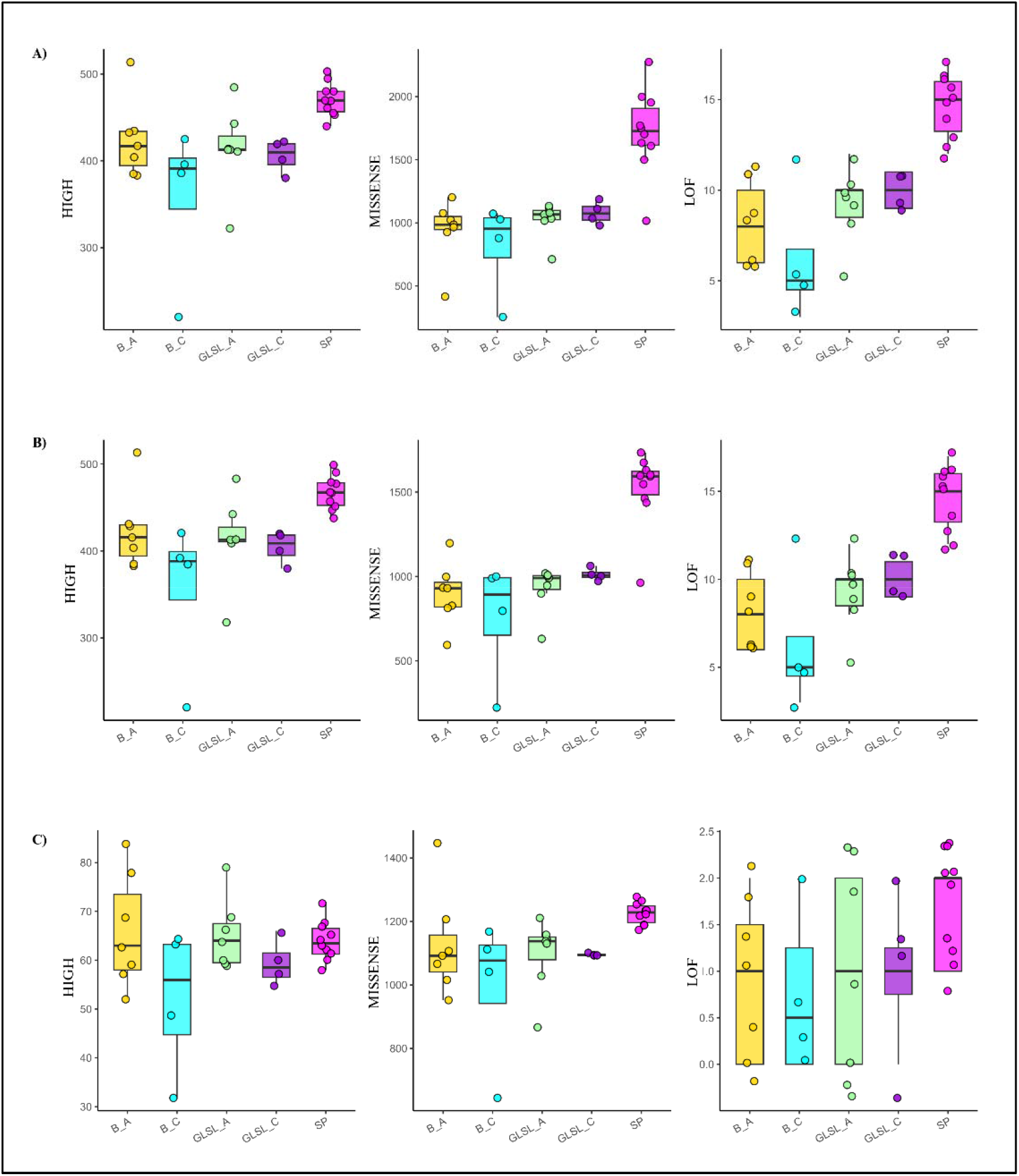
Distribution of predicted functional effects of single nucleotide polymorphism (SNPs) across black bear populations using SnpEff. **a)** Average of all SNPs identified using ANGSD (ROHs and SVs masked). **b)** Average of SNPs overlapping runs of homozygosity **c)** Average of SNPs overlapping structural variants. Boxplots show the number of SNPs classified by SnpEff as HIGH, MISSENSE, or loss of function (LOF) for individuals within each population.

Using the criteria of fixed SVs and ROHs, we identified numerous impacted genes that differed in number among populations. The isolated SP population contained 11 genes associated with fixed SV regions and 29 genes associated with ROH-overlapping regions. GLSL_A contained 26 SV associated genes and one ROH associated gene. B_A contained 30 SV associated genes and seven ROH associated genes. GLSL_C contained 23 SV associated genes and nine ROH associated genes, while B_C contained six SV associated genes and three ROH associated genes. Although each population contained unique genes, several common functional themes were observed. Genes involved in nervous system development, neuronal function, and synaptic processes were identified across multiple populations, including TENM4, RBFOX1, UNC5C, SEZ6L, and SLITRK4. Genes associated with intracellular signaling pathways, including components of the MAP kinase, PI3K, and Wnt signaling pathways, were also identified across populations. Additional genes were associated with cytoskeletal organization and cell adhesion, whereas olfactory receptor genes were identified in both the GLSL_A and GLSL_C populations. The SP population contained genes associated with neuronal signaling and intracellular signal transduction (Table S2, Table S3).

## 4. Discussion

### Population Genomic Consequences of Isolation in Ontario Black Bears

Population isolation reduces gene flow and increases the influence of genetic drift, often leading to increased inbreeding and the loss of genetic diversity (Pelletier et al., 2017; Clendenin et al., 2025; Smeds et al., 2024). Consistent with these expectations, black bears from Saugeen Peninsula (SP) were genetically distinct from mainland Ontario populations and formed a separate genetic cluster in both population structure and admixture analyses (Figure S1a, Figure S2a). In contrast, mainland populations exhibited relatively weak genetic differentiation and extensive shared ancestry, suggesting a history of connectivity and gene flow. Although SP and mainland populations presumably share a common ancestral origin, geographic isolation and increasing urbanization appears to have reduced gene flow between them over time. While a small number of SP individuals showed evidence of recent mainland ancestry, gene flow appears to have been limited. This reduction in connectivity, followed by the effects of genetic drift in a relatively small and isolated population, likely led to the genetic differentiation observed between SP and mainland populations (Pelletier et al., 2017; Clendenin et al., 2025).

The effects of isolation were evident in increased genome-wide homozygosity, a greater abundance of ROHs, and elevated genetic load. Similar genomic signatures have been reported in isolated wildlife populations where reduced connectivity and prolonged demographic isolation have resulted in inbreeding and reduced heterozygosity (Kardos et al., 2023; Cars and Shafer, 2026;). Furthermore, ROHs were more frequently fixed among individuals within the SP population, with SP having the largest number of ROHs suggesting that homozygous genomic regions are likely due to a founder effect given their lengths were not longer than mainland bears (Table 5). Such patterns suggest that the SP bear population may have experienced long-term isolation from mainland populations. Increased homozygosity likely has increased over generations through genetic drift, possibly accelerated by population declines. However, the lack of long ROH, which are associated with more recent inbreeding, suggests that recent mating may not be a caused to observed homozygosity. Instead, the abundance of shorter ROH is more consistent with the lasting genetic effects of historical isolation and bottlenecks (Shafer and Kardos, 2025).

**Table 5.** Summary of fixed runs of homozygosity among Ontario American black bear populations. The number of fixed ROHs, average ROH length, and median ROH length are reported for each population, including SP analyzed with mainland individuals.

| **Population** | **# ROH** | **AVG size in kbp** | **Med Size in kbp** |
|---|---|---|---|
| **SP** | 17 | 59 | 23 |
| **B_A** | 3 | 38 | 46 |
| **B_C** | 3 | 32 | 24 |
| **GLSL_A** | 2 | 178 | 178 |
| **GLSL_C** | 9 | 263 | 118 |

The combination of extensive ROHs and elevated genetic load observed in SP black bears therefore suggests that restricted gene flow and persistently small effective population size have jointly influenced the genomic health of this population. Similar relationships among reduced effective population size, inbreeding, and declines in heterozygosity have been reported across numerous isolated vertebrate populations (Kardos et al., 2023; Cars et al., 2024). The elevated abundance of ROHs also has implications for genetic load (Figure 3a, Figure S3). In small and isolated populations, inbreeding increases the probability that deleterious recessive alleles occur in a homozygous state and are therefore expressed phenotypically (Bosse & van Loon, 2022). The increased number of potentially deleterious variations within ROHs may have consequences for the genomic health of the SP population, as increased homozygosity will expose recessive deleterious alleles and potentially reduce fitness. The elevated ROH abundance in SP, together with the genetic load results, suggests that extended isolation has contributed to reduced genetic diversity and increased expression of deleterious variation, potentially increasing the SP population’s vulnerability to genomic erosion (e.g. Bosse & van Loon, 2022).

The Tajima’s D results provided additional evidence for the demographic consequences of isolation as it was positive (i.e. reduction in rare alleles) in the SP population (Table 1), but negative elsewhere. When considered alongside all other metrics, this result is consistent with a demographic history characterized by limited gene flow and strong genetic drift (Pelletier et al., 2017; Puckett & Clendenin, 2026). Importantly, these findings suggest that the genetic distinctiveness of SP black bears is more likely the result of demographic processes than adaptive divergence. Comparable conclusions have been reported for Louisiana black bears, where population differentiation was primarily associated with bottlenecks, drift, and reductions in genetic diversity rather than adaptive evolution (Puckett & Clendenin, 2026).

Genome-wide diversity is increasingly recognized as a fundamental conservation objective because genomic erosion can limit the ability of populations to respond to future selective pressures (Wold et al., 2021). Maintaining and possibly restoring connectivity may help reduce the long-term genomic consequences of isolation and promote the persistence of black bear populations on the peninsula.

### Structural Variation, Genomic Health, and Evolutionary Implications

Structural variants provided insights into genomic variation that were not fully captured by traditional SNP-based population genomic analyses (Table 2, Table 4, Table 6) and can reveal additional dimensions of genomic variation that may contribute to genomic health and evolutionary potential (Mérot et al, 2020). Structural variants represent an important source of diversity because they can affect large genomic regions containing multiple genes and regulatory elements and thus might be important to conservation (Wold et al., 2021; Stuart et al., 2025). The relationship between SV and diversity was most evident in the SP population (Table 6). Individuals from this isolated population exhibited extensive ROH and elevated genome-wide homozygosity, and many SVs overlapped these homozygous regions (Table 2). Because ROHs represent genomic segments inherited identically by descent, SVs occurring within these regions are more likely to occur in a homozygous state and therefore have a greater opportunity to influence phenotype and fitness. Similar patterns have been reported in Scandinavian wolves, where inbreeding increased the realized SV load by exposing variants that were previously masked in heterozygous individuals (Smeds et al., 2024). Indeed, while some metrics reported here are impacted by sample size, our bootstrapped assessment of load and genomic feature, as well as number of fixed ROHs, suggest increased load in SPs is indeed real (Figure S3).

**Table 6.** Summary of structural variant and run of homozygosity overlaps among Ontario American black bear individuals by population. The total number of genomic overlaps between SVs and ROHs is reported for each population. (B_A, GLSL_A and SP have long read data.)

| **Population** | **Number of SV and ROH overlap** | **Avg. per individual** |
|---|---|---|
| **SP** | 12875 | 1288 |
| **B_A** | 8750 | 1750 |
| **B_C** | 739 | 185 |
| **GLSL_A** | 4806 | 687 |
| **GLSL_C** | 1310 | 328 |

Several broad functional categories were repeatedly associated with genes linked to fixed SVs. These included neuronal development, intracellular signaling, cytoskeletal organization, and cell adhesion. The recurrence of these biological functions across populations suggests that SVs frequently affect genes involved in fundamental cellular and regulatory processes, and at least in Ontario, have range-wide effects, more consistent with species evolution than local adaptation. The candidate genes identified in this study should be viewed as hypotheses for future investigation rather than evidence of functional significance. The chromosomal inversions identified among Ontario black bear populations also warrant cautious interpretation. Chromosomal inversions can have diverse evolutionary consequences depending on their frequency, genomic content, and population context. In some systems, inversions contribute to adaptation by preserving favorable combinations of alleles and reducing recombination among locally adapted genomic regions (Schneller et al., 2025). However, inversions may lead to the accumulation of recessive deleterious variation and increase the genomic consequences of inbreeding. The presence of inversions within Ontario black bears therefore cannot be interpreted as evidence of either adaptation or maladaptation. Instead, these variants represent promising targets for future research aimed at understanding their contribution to population differentiation, genomic health, and evolutionary processes.

## 5. Conclusion

Structural variant analyses support the broader interpretation that demographic history has been a major determinant of genomic variation in Ontario black bears. The relationship between SVs and genetic load in SP bears is noteworthy, where the buildup of SVs could in theory lead to incompatibility (Frankham et al, 2011), potentially limiting natural or anthropogenic attempts at a genetic rescue. However, the biological significance of individual SVs remains uncertain, and future work integrating LR sequencing, functional genomic analyses, and direct assessments of fitness will be necessary to determine their evolutionary and conservation relevance. American black bears from the Saugeen Peninsula showed clear genetic differentiation from mainland populations, elevated ROH, and high genetic load. Our analyses is consistent with drift-driven divergence resulting in SP divergence, not adaptation. Restoring connectivity between SP and mainland populations (genetic rescue) may help reduce loss of genetic diversity. As conservation genomics increasingly moves beyond SNP-based measures of diversity, SVs may provide an important complementary perspective on genomic health and the long-term consequences of population isolation (Wold et al., 2021; Stuart et al., 2025).

## Acknowledgements

We thank the numerous Ontario Ministry of Natural Resources employees who collected samples and the hunters who submitted teeth. We thank K. Beauclerc for DNA extraction, data curation and assistance with sample selection.

## Data availability

The code supporting this work is available in our lab’s GitLab repository: https://gitlab.com/WiDGeT_TrentU/graduate_theses/-/tree/master/AzimiChetabi/Chapter%203?ref_type=heads

## Author Contribution

Based on CRediT(https://credit.niso.org/):

Faezeh Azimi Chetabi: Analysis, Writing – original draft, Writing – review & editing.

Aaron B.A Shafer: Supervision, Writing – review & editing.

Emily E. Puckett: Writing – review & editing.

Joseph M. Northrup: Writing – review & editing.

All authors reviewed and approved the final version of the manuscript.

## Funding

We were supported by Trent University’s International Graduate Scholarship, an Ontario Graduate Scholarship, and Dean’s PhD Scholarship. This work was supported by the Ontario Ministry of Natural Resources, the Natural Sciences and Engineering Research Council of Canada Discovery Grant (grant number: RGPIN-2017-03934), ComputeCanada Resources for Research Groups (grant number: RRG gme-665-ab), Canada Foundation for Innovation: John R. Evans Leaders Fund, and Ontario Early Researcher Award (grant number: #36905).

## Conflict of interest statement

The authors declare no competing interests.

## Notes

### Competing Interest Statement

The authors have declared no competing interest.

