## Supplemental Data for "Measuring American black bear genetic load in fixed structural variants and runs of homozygosity"

**Supplementary Material**

**A)**

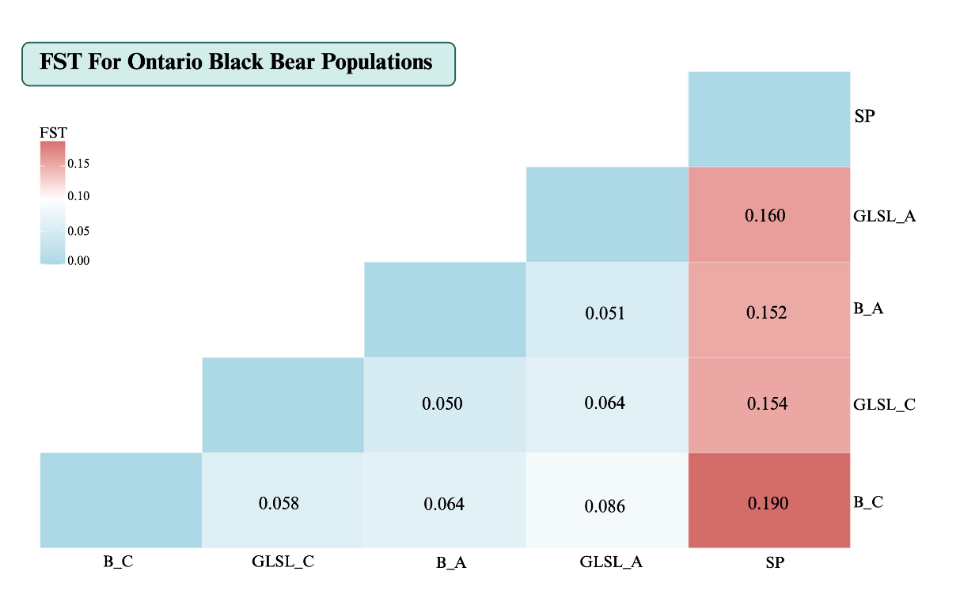

**B)**

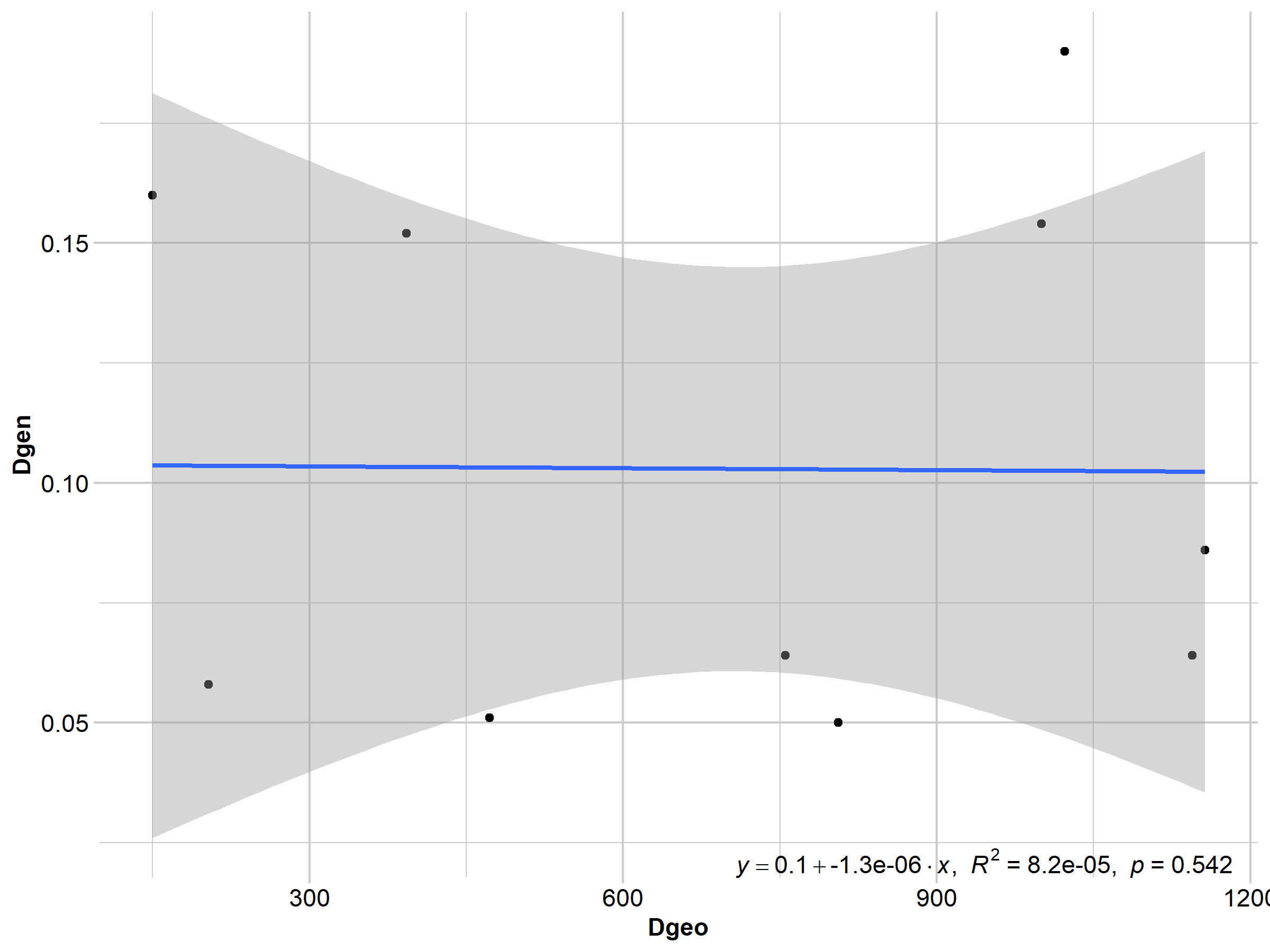

**Figure S1. a)** Pairwise genetic differentiation among Ontario American black bear populations. The heatmap shows pairwise $F_{st}$ estimates between populations, with higher values indicating greater genetic differentiation. **b)** Isolation by distance (IBD) among populations. Each point represents a pairwise comparison between populations, with geographic distance on the x-axis and genetic differentiation ($F_{st}$) on the y-axis. The blue line represents the linear regression, and the gray shaded area indicates the 95% confidence interval.

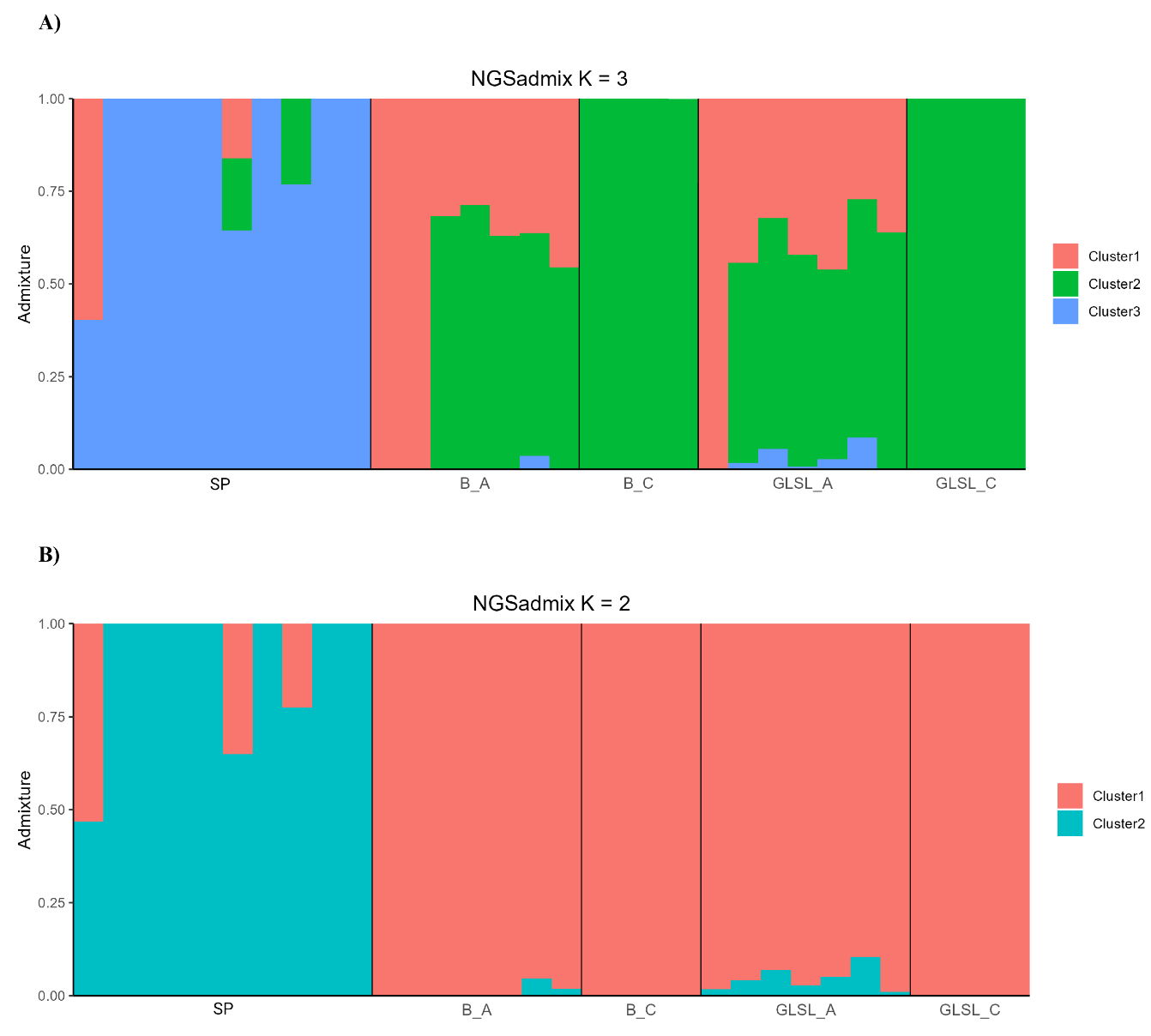

**Figure S2.** Individual ancestry proportions among Ontario American black bear populations estimated using NGSadmix, assuming **a)** three ancestral clusters (K = 3) and **b)** two ancestral clusters (K = 2). Each vertical bar represents an individual, with colors indicating the estimated proportion of ancestry assigned to each genetic cluster. Individuals are grouped by population, with black vertical lines separating populations.

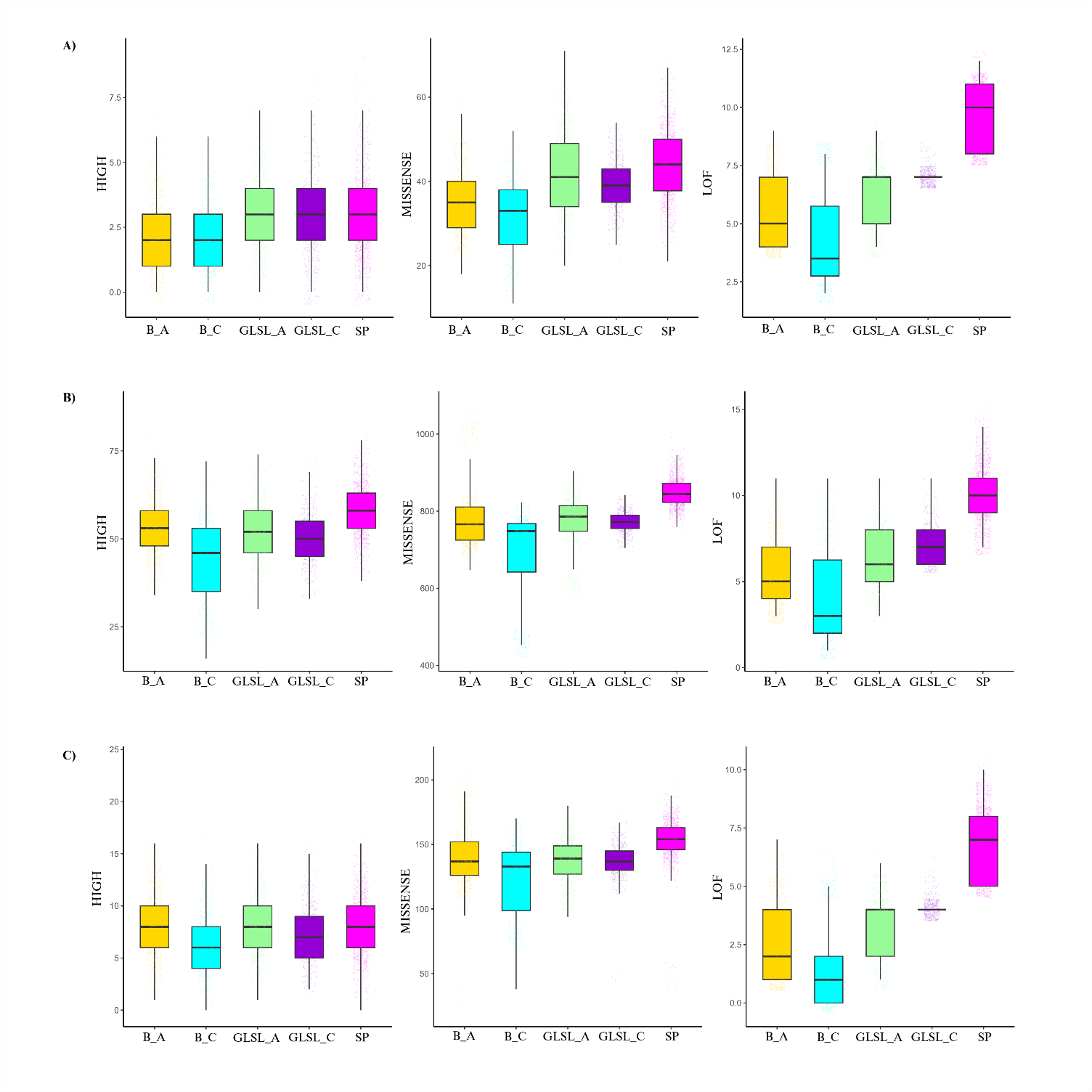

**Figure S3.** Distribution of predicted functional effects of single nucleotide polymorphism (SNPs) across black bear populations using SnpEff (100 files of 1,000,000 sites, containing masked and realized load), Y axis shows counts of SNPs. **a)** All SNPs identified using ANGSD (ROHs and SVs masked). **b)** SNPs overlapping runs of homozygosity **c)** SNPs overlapping structural variants. Boxplots show the number of SNPs classified by SnpEff as HIGH, MISSENSE, or loss of function (LOF) for individuals within each population.

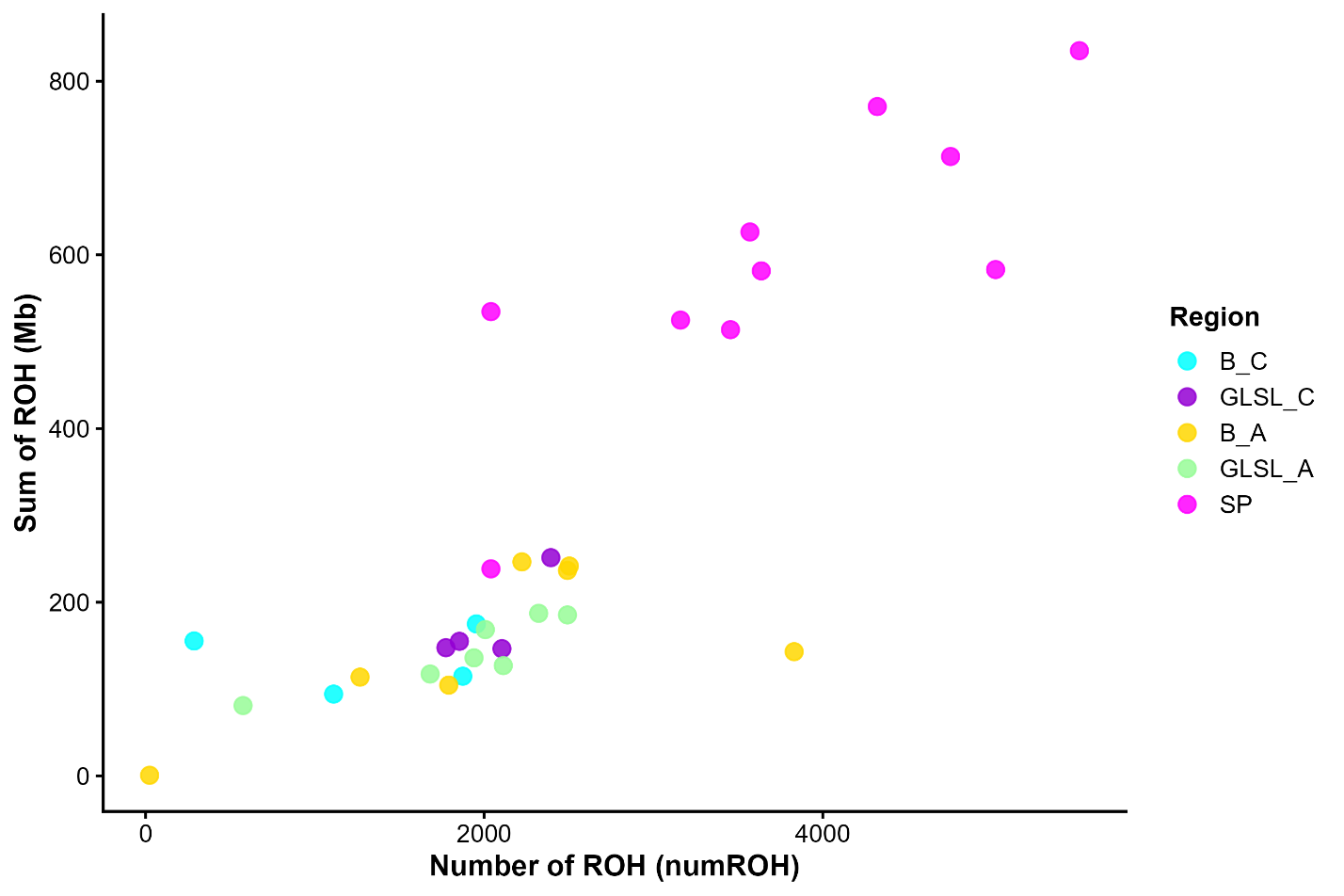

**Figure S4.** Distribution of runs of homozygosity (ROH) across individuals from each region. Each point represents an individual, with the total number of ROH (numROH) on the x-axis and the total length of ROH (sumROH, Mb) on the y-axis.

**Table S1.** Sample information and sequencing characteristics of American black bears used in this study. The table presents sample identifiers, geographic regions, sequencing type (SR: short-read, LR: Long-read), average sequencing coverage depth, breadth of coverage at 1× and 5× and geographic coordinates (latitude and longitude) for each sample. (KFN: Kashechewan First Nation)

| Sample | Region | Type | Coverage depth | Breadth x1 | Breadth x5 | Latitude | Longtitude |
| --- | --- | --- | --- | --- | --- | --- | --- |
| BB0014051 | B_C | SR | 9.91058 | 2491915041 | 2329664170 | 50.63695 | -93.2570545 |
| BB0014056 | B_C | SR | 10.6809 | 2502812024 | 2344490714 | 50.63695 | -93.2570545 |
| BB0014064_S3 | B_C | SR | 8.89912 | 2491874417 | 2211157057 | 50.63695 | -93.2570545 |
| BB0014065_S4 | B_C | SR | 1.66355 | 1546240186 | 21075832 | 50.63695 | -93.2570545 |
| BB0014147_S5 | GLSL_C | SR | 9.24948 | 2498954188 | 2258138454 | 48.87131 | -93.9828688 |
| BB0014176_S6 | B_A | SR | 12.1829 | 2494230824 | 2411625101 | 49.20361 | -82.9460946 |
| BB0014177_S7 | B_A | SR | 11.155 | 2499110128 | 2356443675 | 49.20361 | -82.9460946 |
| BB0014306_S8 | GLSL_C | SR | 10.021 | 2501868094 | 2312732046 | 48.87131 | -93.9828688 |
| BB0014469_S9 | B_A | SR | 11.3615 | 2505834765 | 2377310418 | 49.20361 | -82.9460946 |
| BB0014472_S10 | B_A | SR | 9.61264 | 2500539723 | 2220227219 | 49.20361 | -82.9460946 |
| BB0014527_S11 | GLSL_A | SR | 13.2169 | 2505023352 | 2415270203 | 45.61991 | -79.5854005 |
| BB0014609_S12 | GLSL_A | SR | 9.21871 | 2500915444 | 2251543445 | 45.61991 | -79.5854005 |
| BB0014619_S13 | GLSL_A | SR | 9.81997 | 2496816276 | 2293401016 | 45.61991 | -79.5854005 |
| BB0015636_S14 | GLSL_C | SR | 10.2633 | 2490111676 | 2347189018 | 48.87131 | -93.9828688 |
| BB0037469_S15 | GLSL_C | SR | 11.0258 | 2500495305 | 2349848278 | 48.87131 | -93.9828688 |
| BB0037677_S16 | GLSL_A | SR | 10.6953 | 2490571512 | 2365361169 | 45.61991 | -79.5854005 |
| BB0050853_S17 | SP | SR | 9.8498 | 2498596967 | 2298564154 | 45.81534 | -81.4906309 |
| BB0059320_S26 | SP | SR | 10.9282 | 2500642760 | 2352959336 | 45.81534 | -81.4906309 |
| BB0059328_S25 | SP | SR | 8.45743 | 2497721064 | 2178450468 | 45.81534 | -81.4906309 |
| BB0059559_S19 | SP | SR | 7.7181 | 2495259281 | 2080749225 | 45.81534 | -81.4906309 |
| BB0059560_S20 | SP | SR | 8.19053 | 2487456957 | 2163911445 | 45.81534 | -81.4906309 |
| BB0059561_S22 | SP | SR | 8.74675 | 2486524733 | 2232426185 | 45.81534 | -81.4906309 |
| BB0059793_S24 | SP | SR | 8.75593 | 2494884414 | 2205455578 | 45.81534 | -81.4906309 |
| BB0059794_S27 | SP | SR | 10.9644 | 2496887301 | 2382941409 | 45.81534 | -81.4906309 |
| BB0059799_S21 | SP | SR | 10.1508 | 2498263849 | 2311388640 | 45.81534 | -81.4906309 |
| BB0059800_S23 | SP | SR | 3.26879 | 2336376793 | 497600019 | 45.81534 | -81.4906309 |
| BB0059803_S18 | SP | SR | 5.95451 | 2475927284 | 1641900772 | 45.81534 | -81.4906309 |
| BB03_S88_L00 | Peawanuck | SR | 10.2177 | 2499362337 | 2180557938 | 54.9939 | -85.4277 |
| bc2045_250228_160914 | KFN | LR | 15.2328 | 2504785988 | 2459932125 | 52.2915 | -81.6407 |
| m84090_230801_152430 | SP | LR | 22.603 | 2504315399 | 2480870348 | 44.569 | -80.9406 |
| m84090_240118_205717 | B_A | LR | 30.3804 | 2507787045 | 2488306849 | 47.6307 | -80.87418 |
| m84255_250228_160914 | GLSL_A | LR | 14.9048 | 2490985571 | 2444207562 | 45.19996 | -78.243267 |

**Table S2.** Genes associated with fixed structural variants (SV) in the study population. Each gene is associated with an SV identified as fixed within the population.

| **Population** | **Gene name** | **Gene ID** | **Function** |
| --- | --- | --- | --- |
| B_C | NDST3 | 9348 | involved in heparan sulfate proteoglycan biosynthetic process |
|  | CST3 | 100462660 | enables cysteine-type endopeptidase inhibitor activity |
|  | CST3 | 100462660 | enables cysteine-type endopeptidase inhibitor activity |
|  | HK3 | 510616 | involved in glucose metabolic process |
|  | APC | 324 | involved in negative regulation of cell population proliferation |
|  | SLC35F3 | 148641 | enables choline transmembrane transporter activity |
| GLSL_C | SLIT3 | 6586 | involved in embryonic olfactory bulb interneuron precursor migration |
|  | PDCD6IP | 10015 | involved in multivesicular body assembly |
|  | TBC1D20 | 128637 | involved in positive regulation by virus of viral protein levels in host cell |
|  | KIZ | 458130 | spindle organization |
|  | HUB1 | 855767 | involved in cell morphogenesis involved in conjugation with cellular fusion |
|  | UNC5D | 137970 | enables netrin receptor activity |
|  | IL-16 | 574100 | lymphocyte chemoattractant factor |
|  | RBFOX1 | 100172936 | involved in regulation of alternative mRNA splicing, via spliceosome |
|  | KIFAP3 | 518402 | involved in microtubule-based movement |
|  | PARP1 | 286764 | involved in double-strand break repair |
|  | FILIP1 | 27145 | involved in modification of postsynaptic structure |
|  | LRRC43 | 254050 | protein-protein interactions |
|  | VPREB1 | 7441 | encodes the iota polypeptide chain that is associated with the Ig-mu chain |
|  | Or13l2 | 258272 | enables odorant binding |
|  | JAZF1 | 221895 | enables transcription corepressor activity |
|  | Cldnd1 | 224250 | enables molecular_function |
|  | FTH1 | 281173 | Iron ion homeostasis and iron storage |
|  | Megf10 | 291445 | enables Notch binding |
|  | HSD17B4 | 3295 | involved in estrogen metabolic process |
|  | CHRM3 | 281685 | - |
|  | CHRM3 | 281685 | enables G protein-coupled acetylcholine receptor activity |
|  | Ube2k | 53323 | enables ubiquitin conjugating enzyme activity |
|  | BPNT2 | 54928 | enables 3',5'-nucleotide bisphosphate phosphatase activity |
| B_A | ADAMTS9 | 537051 | enables metalloendopeptidase activity |
|  | CGA | 280749 | enables follicle-stimulating hormone activity |
|  | FLI1 | 529124 | enables sequence-specific DNA binding |
|  | UTP20 | 27340 | enables protein binding |
|  | Scyl2 | 213326 | enables protein kinase activity |
|  | Rbms1 | 56878 | enables mRNA 3'-UTR binding |
|  | CFAP161 | 100156420 | involved in cilium assembly |
|  | CLRN2 | 645104 | - |
|  | PRICKLE2 | 166336 | - |
|  | Snx2 | 67804 | enables epidermal growth factor receptor binding |
|  | WDR11 | 55717 | involved in head development |
|  | FHIP2A | 57700 | enables protein binding |
|  | Ablim1 | 226251 | enables actin filament binding |
| GLSL_A | ICA1L | 130026 | Associated with intracellular membrane trafficking |
|  | Dnaja3 | 360481 | Molecular chaperone activity and protein folding |
|  | UCHL5 | 51377 | Binding to a proteasome, a large multisubunit protein complex that catalyzes protein degradation |
|  | Actr3b | 362298 | A cellular structure that forms the internal framework of eukaryotic and prokaryotic cells |
|  | DGKB | 1607 | diacylglycerol kinase activity (ATP |
|  | CMSS1 | 616567 | A large ribonucleoprotein complex considered to be the earliest preribosomal complex. |
|  | Tcerg1l | 361669 | transcription cofactor activity |
|  | UNC5C | 8633 | netrin receptor activity involved in negative chemotaxis, repulsive netrin receptor activity |
|  | AGO2 | 27161 | enables RNA endonuclease activity |
|  | FAM110B | 90362 | A structure comprised of a core structure (in most organisms, a pair of centrioles) |
|  | CNTN6 | 27255 | The attachment of an adhesion molecule in one cell to an identical molecule in an adjacent cell |
|  | MYOCD | 93649 | Binding to a sequence-specific DNA binding RNA polymerase II transcription factor |
|  | NDST3 | 9348 | involved in heparan sulfate proteoglycan biosynthetic process |
|  | SLC30A7 | 100156498 | cobalt, zinc uptake permease activity |
|  | CHORDC1 | 101925655 | - |
|  | TBC1D20 | 128637 | involved in positive regulation by virus of viral protein levels in host cell |
|  | RASSF3 | 536808 | The cellular process in which a signal is conveyed to trigger a change in the activity or state of a cell |
|  | TBK1 | 533141 | involved in peptidyl-serine phosphorylation |
|  | rnf7 | 503534 | Binding to a member of the cullin family |
|  | WDR7 | 23335 | ny process that reduces the pH of the vacuole |
|  | SSTR4 | 6754 | somatostatin receptor activity |
|  | Map2k6 | 26399 | Catalysis of the reactions: ATP + protein |
|  | OR5T1 | 390155 | perception of smell, detection of chemical stimulus, perception of smell |
|  | OR5T1 | 390155 | perception of smell, detection of chemical stimulus, perception of smell |
|  | AP5M1 | 55745 | protein amino acid binding, glycoprotein binding |
|  | rps29 | 100304579 | The action of a molecule that contributes to the structural integrity of the ribosome |
| SP | mapk10 | 108711504 | acts_upstream_of_or_within rhythmic process |
|  | AGO2 | 27161 | enables RNA endonuclease activity |
|  | OPRK1 | 4986 | involved in immune response |
|  | DOCK9 | 23348 | involved in regulation of Rho protein signal transduction |
|  | SLC5A8 | 160728 | involved in nicotinate metabolic process |
|  | TBC1D20 | 128637 | involved in positive regulation by virus of viral protein levels in host cell |
|  | TBK1 | 533141 | involved in peptidyl-serine phosphorylation |
|  | RPL8B | 850682 | involved in maturation of LSU-rRNA |
|  | SSTR4 | 6754 | somatostatin receptor activity |
|  | CELF2 | 10659 | involved in mRNA splice site recognition |
|  | SEZ6L | 23544 | involved in adult locomotory behavior |

**Table S3.** Genes associated with fixed runs of homozygosity (ROH) in the study population. Each gene is associated with an ROH identified as fixed within the population.

| **Population** | **Gene name** | **Gene ID** | **Function** |
| --- | --- | --- | --- |
| B_C | Slitrk4 | 245446 | protein amino acid binding, glycoprotein binding |
|  | AMOT | 154796 | Involved in angiogenesis, cell migration, and cell junction organization |
|  | DMD | 606758 | protein amino acid binding, glycoprotein binding |
| GLSL_C | MAGEA6 | 4105 | Involved in regulation of protein ubiquitination and cellular signaling |
|  | ZNF41 | 7592 | protein amino acid binding, glycoprotein binding |
|  | RPS6KA3 | 6197 | Mg binding, magnesium binding |
|  | APOBEC3Z3 | 100127022 | (deoxy)cytidine deaminase activity, cytidine aminohydrolase activity, |
|  | FRS2 | 10818 | eurotrophin TRKA receptor ligand |
|  | C23H9orf72 | 100053503 | Involved in intracellular membrane trafficking and autophagy |
|  | CD163 | 477704 | ransmembrane, integral component of membrane |
|  | CLEC4E | 26253 | Binding to a carbohydrate, which includes monosaccharides |
|  | SPATA31D1 | 389763 | Predicted to be involved in spermatogenesis |
| B_A | NM_173694.5 | 286410 | ATPase-coupled phospholipid transporter activity |
|  | Gpc3 | 25236 | Involved in cell growth regulation and embryonic |
|  | MAGEA6 | 4105 | Involved in regulation of protein ubiquitination and cellular signaling |
| GLSL_A | CHD6 | 524776 | ATP hydrolysis-dependent chromatin remodeler activity |
|  | FGF4 | 618474 | animal organ morphogenesis |
| SP | Hapstr2 | 279618 | Binding to a ubiquitin protein ligase enzyme |
|  | CXorf66 | 347487 | protein amino acid binding, glycoprotein binding |
|  | Fgf13 | 14168 | The function that stimulates a cell to grow or proliferate |
|  | PPP1R2C | 80316 | hosphoprotein phosphatase inhibitor activity |
|  | PPP1R2C | 80316 | hosphoprotein phosphatase inhibitor activity |
|  | BCOR | 784529 | transcription co-repressor activity, RNA polymerase II transcription co-repressor activity |
|  | BCOR | 784529 | transcription co-repressor activity, RNA polymerase II transcription co-repressor activity |
|  | H1-10 | 8971 | Binding to the DNA portion of a nucleosome |
|  | EFCC1 | 617677 | transcription co-activator activity |
|  | IQSEC1 | 9922 | GDP-dissociation stimulator activity |
|  | IQSEC1 | 9922 | GDP-dissociation stimulator activity |
|  | Hdac11 | 232232 | Removal of an acetyl group from a lysine residue in a histone |
|  | Hdac11 | 232232 | Removal of an acetyl group from a lysine residue in a histone |
|  | TMEM38B | 615646 | isoform-specific homophilic binding |
|  | TAL2 | 6887 | protein dimerization activity |
|  | FKTN | 2218 | Catalysis of the transfer of a substituted phosphate group |
|  | Rbms3 | 207181 | adenylate/uridylate-rich element binding |
